# CurveKinetix: A user-friendly application for analysis of biological time series data

**DOI:** 10.64898/2026.11.24.768863

**Authors:** Matthew Warren, Alyson Hockenberry

**Affiliations:** Loyola University Chicago

## Abstract

Of interest to many biologists is how organisms change over time: their growth, their response to a stimulus, their movement. How quickly a process starts, accelerates, slows, or changes direction, often carries critical biological information for hypothesis testing. Extraction of these parameters, however, requires computational expertise or reliance on existing software not designed for the needed analysis. We present here CurveKinetix, a graphical user interface application that helps researchers extract quantitative parameters from time series data without writing code. Users can quickly go from instrument data files to estimation of parameters using three complementary methods to visualized data customization and export.

We exhibit CurveKinetix’s utility in two case studies. In the first, we show expression of an operon correlates with slowing of population level growth. In the second, we enumerate viral fusion dynamics, demonstrating CurveKinetix’s framework can be applied to a range of biological questions.

By making flexible quantitative analysis straightforward, CurveKinetix helps experimentalists extract more information from their data and supports clearer, more rigorous interpretation of time-resolved experiments.

**Significance:** We developed CurveKinetix to make quantitative analysis of biological time series data accessible to researchers who do not write code. It can process both simple and complex curves, helping users extract meaningful parameters from their data to improve the interpretability and rigor of their time-resolved experiments.

## Introduction

Biological processes happen across timescales. While endpoint measurements of absolute values like biological yield, gene expression, or protein levels are sufficient for testing some hypotheses, others require information that exists in how a given value changes over time. Extracting these values from time-resolved datasets generally requires *curve fitting*, the process of finding the mathematical equation that best describes the shape of the data. Once the user has this equation, they can then infer the value of interest from their dataset.

Automated data acquisition platforms like plate readers and microscopes have made the gathering of high-resolution time series data routine for many biologists. Extraction of the pertinent values from these datasets, however, requires quite some coding fluency or reliance upon software that fit boilerplate mathematical functions. The result is that many datasets are summarized by values that miss subtle, meaningful features, leaving important data untapped.

For a concrete example, bacterial population growth characteristics have long been approximated by assuming they grow in a sigmoidal or S-shaped curve^1^. To estimate the population’s lag time, growth rate, and carrying capacity, the optical density of a culture is measured over time, and often evaluated in one of two ways. The first approach is commonly to fit a growth curve model, such as the logistic model, to the measurements. The second common approach is to log transform the measurements and fit a straight line to the exponential growth phase. From the fitted model these values of interest can be estimated.

The assumption that the growth follows a sigmoidal curve holds in some cases, making the estimation of these values using these methods perfectly acceptable. Growth of non-model organisms, growth of populations during metabolic shifts, and measuring the impact of phage on bacterial populations, however, are just a few examples in which the growth curves do not follow a sigmoidal pattern. Existing tools (Table 1) typically offer a set of curve fitting options that work well when the populations follow a sigmoidal growth pattern but are insufficient to describe more complex trajectories. Existing tools that are more appropriate for non-sigmoidal curves often require a higher level of coding fluency, making rigorous quantitative analysis of these data out of reach for many biologists.

**Table 1.** Comparison of existing tools for time series data analysis.

| SOFTWARE | ACCESS | DATA INPUT & PREPARATION | ANALYSIS METHODS | EXPERIMENTAL GROUPING | PLOTTING | FIGURE FORMATS | DATA & WORKFLOW EXPORT |
|---|---|---|---|---|---|---|---|
| CURVEKINETIX | GUI; local (R) | Copy/paste; file/layout import; column mapping | Sigmoid models; derivatives; breakpoint regression | Metadata; biological sample IDs; technical curves | Curves; parameters; facets; appearance controls | PNG, PDF, SVG | CSV; projects; run records |
| GCPLYR<sup>3</sup> | R package | Block/wide/tidy; data reshaping | Derivatives; lag; density; AUC; diauxie | Design files; metadata joins | External ggplot2 workflow | Via external R plotting | R data frames; scripted export |
| QURVE<sup>11</sup> | GUI; local (R) | Reader parsers; custom/tidy tables; sample maps | Sigmoid models; log-linear; splines; biphasic | Samples; replicates; concentrations | Curves; parameters; grids; figure controls | PNG, PDF | CSV; RData; PDF/HTML reports |
| GROWTHCURVER<sup>12</sup> | R package | Wide time/well table; numeric values | Logistic fitting; model/empirical AUC | Per-well IDs; grouping in user code | Built-in curve/fit plots | PDF; R graphics | Fit objects; per-well result tables |
| GROWTHRATES | R package | Prepared long-format data frames | Nonlinear/ODE; log-linear; smoothing splines | Formula-based subsets; replicate identifiers | Built-in fit plots; external ggplot2 | Via R graphics | Fit objects; extracted result tables |
| KINBIONT<sup>13</sup> | Julia package | Wide CSV; arrays/DataFrames; annotations | Nonlinear/ODE; log-linear; derivatives; segmented fitting | Replicate averaging; metadata; downstream learning | Scripted Plots.jl; clustering/learning views | Via Julia plotting | CSV results; fits; model comparisons |
| AMIGA<sup>14</sup> | Python; command line | Well/time tables; metadata | GP models; derivatives; diauxie; group tests | Metadata; replicate pooling; covariate tests | Curves; derivatives; plate grids; heatmaps | PDF | Tab-delimited results; predictions; cleaned data |
| GRAPHPAD PRISM | GUI; commercial | Paste/import; XY or grouped tables | Growth models; custom equations; AUC | Replicate columns; groups; shared fit parameters | Curves; parameters; styling; layouts | PNG, PDF, SVG, TIFF, EPS | CSV/TXT/XML; Prism projects |

We developed CurveKinetix to address this issue. CurveKinetix was developed for users with no coding experience so they could reliably estimate pertinent values from their time series data. While originally developed to characterize complex growth patterns of bacterial populations, we noticed the software is useful for a range of biological time series data and can be broadly applied to extract parameters useful in addressing many biological questions.

## Results and Discussion

### The CurveKinetix framework

CurveKinetix’s framework is simple (Fig 1): the user drags their data files into CurveKinetix’s graphical user interface and indicates background subtraction and data fitting preferences, and the software reformats the data, performs the desired transformations and fits, pools the appropriate biological and technical replicates, and presents the results of the analyses. All data are exportable as CSV files for analysis in Excel or R if desired. Projects can be saved with all progress preserved, making it possible to return to an analysis without needing to re-input anything. The user can also systematically explore their data and estimated parameters using CurveKinetix-integrated visualization tools, allowing for checking the goodness of fit of the models as well as direct comparisons of estimated parameters between experimental conditions.

**Fig 1.**
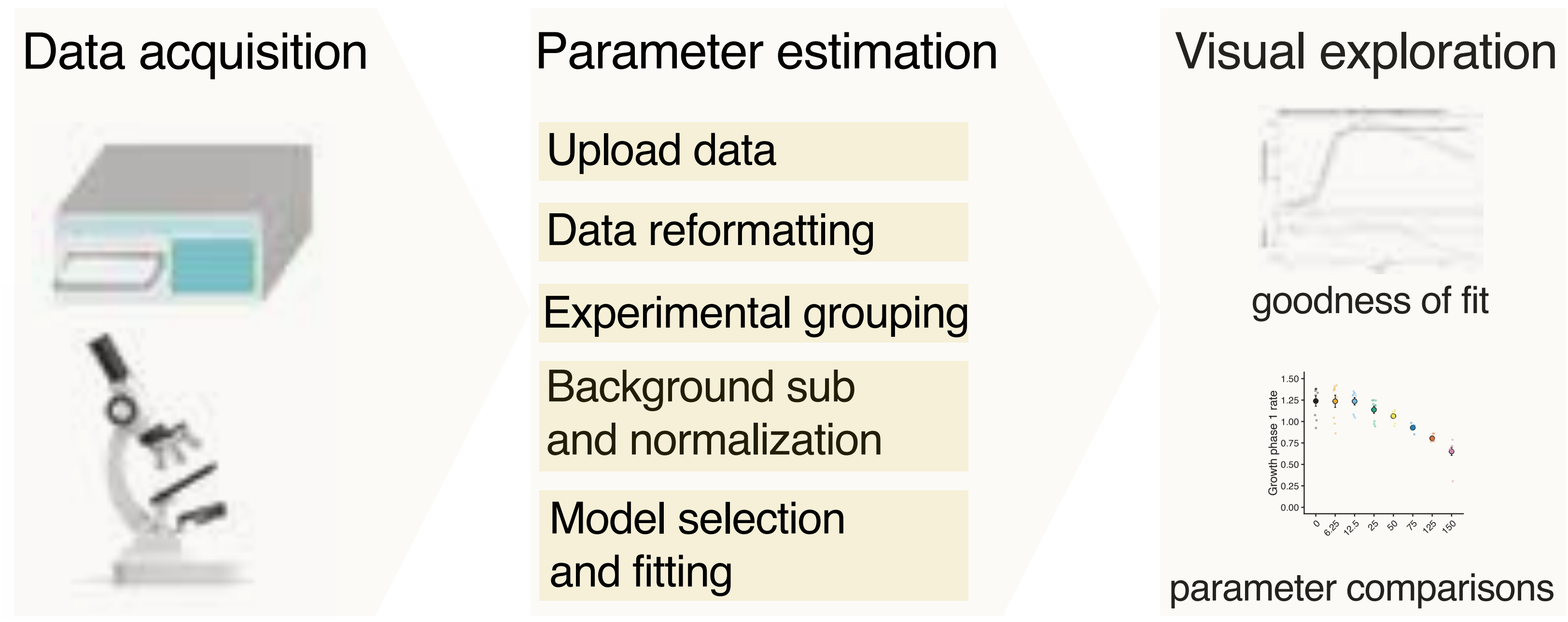
The CurveKinetix Framework. CurveKinetix can analyze time-series data from a range of experimental methods: optical density, colony forming units, fluorescence intensity, enzymatic activity, *etc*. These measurement and time values are then uploaded into CurveKinetix, either by dragging in a file or by copying and pasting the data into the graphical user interface. From there, the user designates preferences for replicate pooling, background subtraction or normalization, and which models to fit. The user can then visually explore their data, examining the raw or processed trajectories over time, the goodness of fit of the models to their data, and a comparison of the extracted parameters over different conditions. Plots are fully customizable and exportable for downstream use.

### Data Preparation

A major pain point in adopting any software is often simply getting data into it in the first place. In CurveKinetix, data preparation is handled by the application. Users can either paste measurement grids and accompanying experimental layouts directly into the Data Wizard or upload supported data and layout files in bulk. With the copy-and-paste method, data does not have to be in a particular data format. Data prepared in the wizard is validated and then passed onto the analysis steps.

### Methods of parameter estimation

CurveKinetix was originally designed to extract plate-reader derived growth curve metrics. Growth of bacterial populations is quantified by measuring the number of individuals over time and is often considered to consist of lag, exponential, and stationary phases (Fig 2). The lag phase is a period of no growth and is attributed to adaptation by the cells to the fresh medium; in many cases of many lab strains, the lag phase is often indetectable. The exponential phase is when the population density increases rapidly. Stationary phase initiates as the cells have exhausted the nutrients in their growth medium and cease dividing. When plotting the number of cells over time, the data can take the shape of an S-shaped or sigmoidal curve; the numbers increase slowly, then quickly, and then slowly again.

**Fig 2.**
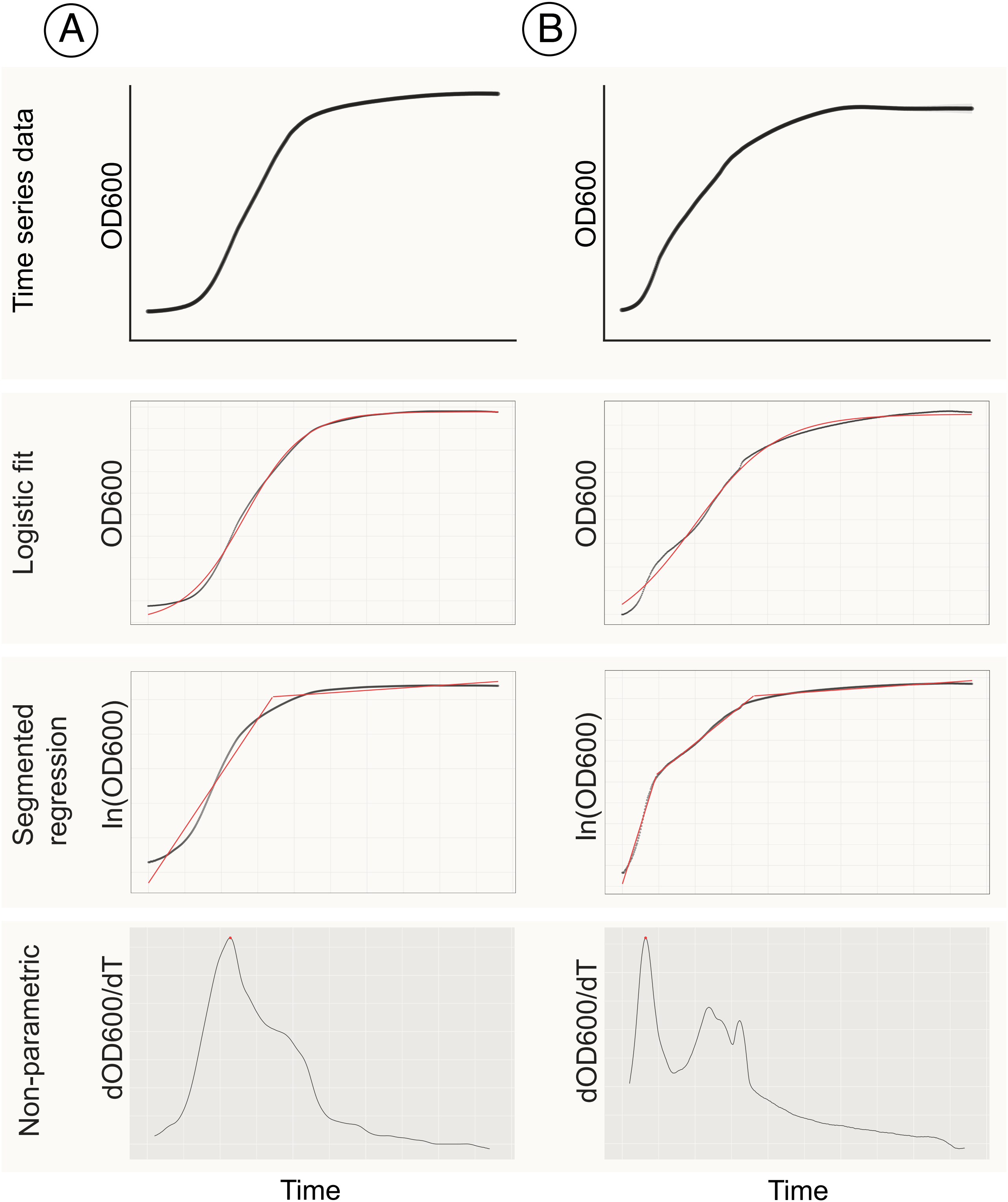
How different models act on different data shapes. *Salmonella* cells can grow in sigmoidal (A) and non-sigmoidal (B) patterns depending on environmental conditions. We fit each of these curves with logistic, segmented regression, and non-parametric models to demonstrate how different curve shapes behave with each method.

There are many cases where bacterial growth does not conform to a sigmoidal curve. A common one is when cells enter a death phase after stationary phase. Another example is diauxic growth, where cells consume a preferred nutrient source and exhaust it before consuming a second nutrient source. In this example, the growth curves take a characteristic double S-shape that is not accurately described by sigmoidal models. In other cases, bacterial growth curves can show changes in growth patterns that may be meaningful to capture but are ultimately lost when estimating parameters using sigmoidal models.

To quantitatively describe the growth patterns of bacterial populations, CurveKinetix includes three modules to fit time-series data. These include parametric fits (Logistic, Gompertz, Gompertz with explicit lag, and Generalized Logistic), break-point analysis (via segmented regression), and non-parametric analysis, which are described in more detail below and in the Supplemental information. Through its creation, we realized that its ability to describe temporal data does not restrict its use to only bacterial growth curves, rather nearly any time series data can be quantitatively analyzed using the application.

### Parametric fitting

For many applications, fitting a curve to growth data is perfectly sufficient to extract useful growth metrics, such as lag time, growth rate, and carrying capacity (see Supplemental information for how these parameters are estimated), which can then be compared to reveal how different nutrient regimes, chemical treatments, or other environmental factors influence growth. While traditionally useful for estimating population growth metrics, fitting a curve to data can also be useful in extracting metrics about gene expression dynamics, enzyme kinetics, and other trajectories that follow similar patterns.

### Breakpoint analysis

Parametric fits can often fail to describe biologically meaningful nuance in time-series data. To address this need, CurveKinetix has a built-in module for breakpoint analysis^2^. In this analysis setting, the data are log-transformed so exponential values becomes linear. A number of lines (the number either chosen automatically or by the user) are fit to the data, such that different phases of the data can be identified, and their rates of change and timepoints in which they start and end can be enumerated. This is useful for quantitatively examining diauxic shifts, death phases, phage dynamics, gene expression data, and other phenomena where the data are not sigmoidal.

### Nonparametric analysis

Lastly, we’ve integrated non-parametric analyses into CurveKinetix^3^. The software will calculate the change in signal over time (first derivative) and identify features in the first derivative that allow for estimation of growth rates, lag time, and more (see Supplemental information for how these parameters are estimated and can be interpreted). This is useful for analyzing data that has characteristics not well captured by parametric models.

### Comparisons, Plotting, and Figure Preparation

Any metadata or annotations that were supplied can be utilized when selecting results, creating comparisons, and constructing plots. Thus, identifiers such as strain, treatment, and experiment id’s can be used to filter and group data or to split figures into panels. This also enables one to identify which technical curves belong to the same biological replicate. With this, technical replicates can be averaged and summaries calculated across biological replicates.

For curve and parameter comparison figures, CurveKinetix allows for a great deal of customization. Users can arrange panels, set custom colors and labels, adjust fonts and legends, and reorder labels. Figures can be exported in PNG, PDF, or SVG formats for seamless insertion into slide decks or manuscripts.

### CurveKinetix applications

#### Case study: correlating bacterial growth patterns and gene expression in different environmental conditions

When *Salmonella* cells grow in LB medium, the growth pattern does not follow a logistic curve. Rather, exponential growth starts at a high rate (phase 1), slows to a lower growth rate (phase 2), and then enters stationary phase. It has been noted that the change in exponential growth rate occurs at roughly the same point as initiation of *Salmonella* pathogenicity island 1 (SPI-1) expression^4^.

Short-chain fatty acids (SCFAs) are shown to reduce SPI-1 expression at the population-level^5–7^, and we hypothesized that because population-level growth and SPI-1 expression are correlated, SCFAs will change how these cells grow. To test this idea, we inoculated a *Salmonella* strain with a chromosomally inserted SPI-1 GFP reporter (SPI1*-gfp)* and its non-fluorescent parent (wt) into LB medium containing increasing physiological levels of SCFAs and measured OD600 and GFP fluorescence values every 3 minutes over a 24 hour period (data adapted from^7^, supplementary file 1). Included in these experiments were uninoculated control wells containing matching concentrations of SCFAs and three technical replicates. The experiments were repeated thrice, yielding 9 measurements for each condition.

We then moved to CurveKinetix for the data analysis (Fig S1). The raw plate reader data were uploaded and reformatted using the built-in wizard. We proceeded to the integrated background subtraction. To remove OD600 background signal, we subtracted the uninoculated well signal from wt and SPI1-*gfp* wells by SCFA concentration. Bacterial autofluorescence in the GFP channel was removed by subtracting the GFP signal of wt wells from the SPI1-*gfp* wells.

A visual inspection of the data indicates SCFAs alter population-level growth patterns and SPI-1 expression (Fig 3A). Commonly used area under the curve (AUC) analyses detect no differences in growth dynamics by SCFA concentration but do show a dose-dependent decrease in SPI-1 expression (Fig 3B).

**Fig 3.**
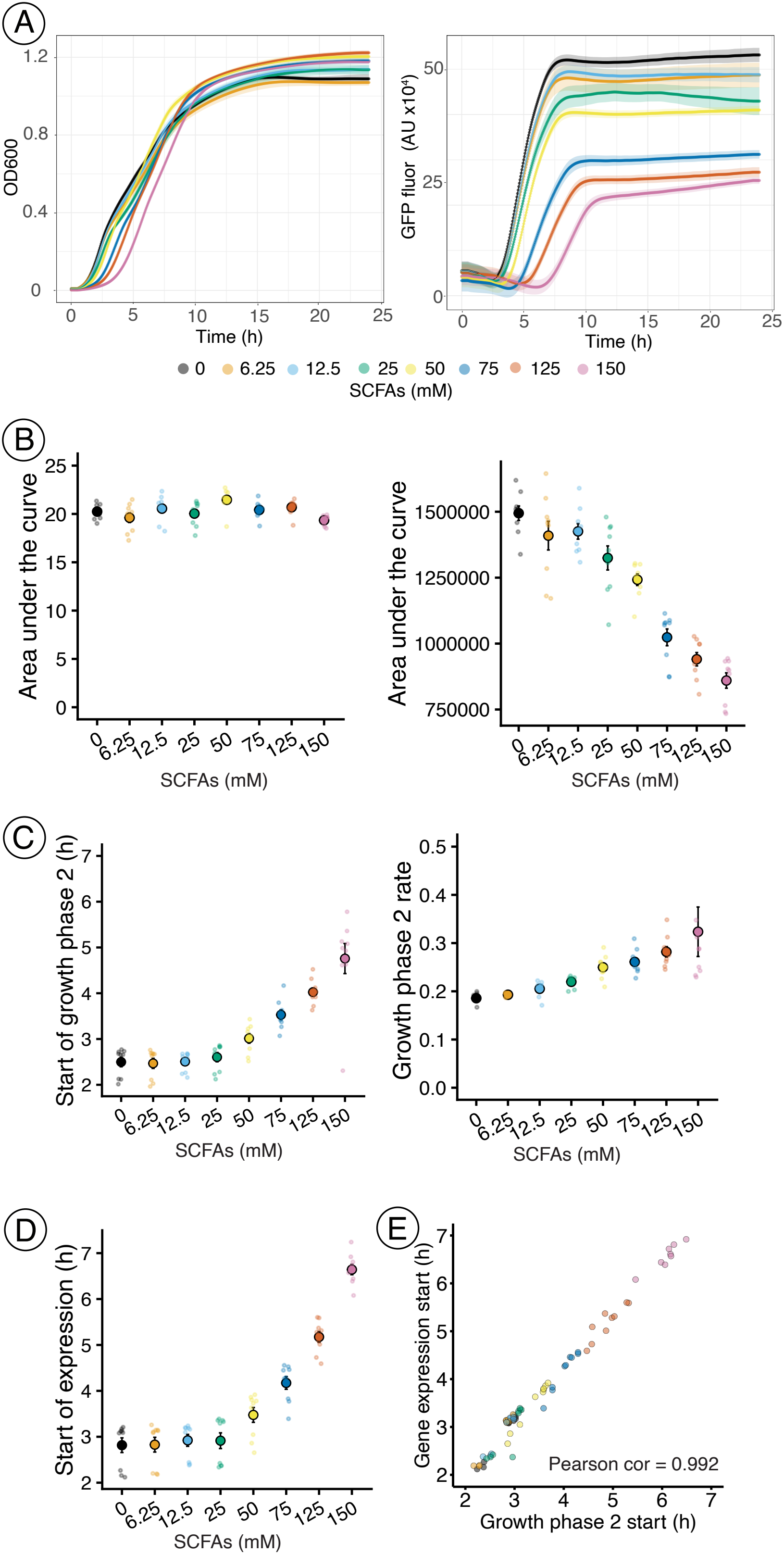
Short-chain fatty acids change *Salmonella enterica* growth and virulence gene dynamics. **A** Optical density (left) and GFP fluorescence (right) values of *Salmonella* SPI-1-gfp cells over 24 hours in the presence of SCFAs at the indicated concentration. Line indicates mean of 3 technical replicates x 3 biological replicates; shaded area is the standard error of the mean (SEM). **B** Area under the curve values for optical density (left) and GFP fluorescence (right) curves. Mean ± SEM. **C** Estimated values of the starting time of the second growth phase (left) and rate of growth in the second growth phase (right) as estimated by CurveKinetix’s breakpoint analysis. Mean ± SEM. **D** CurveKinetix’s estimation of when GFP expression initiated in each sample, as determined by breakpoint analysis. Mean ± SEM. **E** Each sample’s estimated growth phase 2 start time plotted against estimated GFP fluorescence expression start, colored by SCFA treatment condition. Pearson’s correlation = 0.992.

We then shifted to a breakpoint analysis to identify how growth changes in the different conditions (Fig 3C, Fig S1). The breakpoint analysis identified 3 growth phases, with the second visually correlating with SPI1 expression. SCFA treatment delayed the start of growth phase 2 and increased the growth rate during that phase. This is consistent with observations showing SCFAs reduce SPI-1 expression and SPI-1 expression slows growth rates^4,7^.The breakpoint analysis of the GFP fluorescence data similarly identified 3 phases and shows that SCFAs delay the time of initiation of SPI-1 expression (Fig 3D). A deeper analysis shows that the start of growth phase 2 is highly correlated with the start of SPI-1 expression (Fig 3E, *Pearson correlation = 0*.*992*). The accumulation of GFP fluorescence initiates about 10 minutes after the initiation of growth phase 2 (*10*.*56 ± 1*.*53 minutes, mean ± SEM*), which can be attributed to the time of maturation of the GFP (*gfpmut2*, maturation half time (t_50_) ∼6-7 minutes, t_90_ ∼29 minutes^8^).

In summary, SCFAs change both population-level growth and SPI-1 expression kinetics. While the changes in the growth dynamics were not captured by the typical area under the curve analysis, the breakpoint analysis shows that SCFAs delay the start of growth phase 2 and increase the population-level growth during that window. While traditionally used AUC of the GFP data indicated SCFAs reduce SPI-1 expression, this analysis is unable to reveal that these compounds also delay the onset of SPI-1 expression. Furthermore, combining the break-point analysis data substantiated the idea that the slower growth phase is correlated with SPI-1 expression, even in the presence of compounds that modulate the timing and intensity of SPI-1 expression. CurveKinetix allowed us to quickly and easily extract meaningful information that is missed by commonly used analyses.

### Case study: comparing viral fusion kinetics

Viral infection is initiated by particle entry into a host cell. For SARS-CoV-2, the spike protein is a major determinant of this step of infection, and consistent with its function in host cell entry, the spike protein is under heavy evolutionary selection. As SARS-CoV-2 evolved during the pandemic, a spike protein variant bearing a H655Y mutation arose and became fixed in the population^9^. This mutation is associated with slowing viral fusion, which stabilizes particle adsorption and ultimately enhances infection^10^.

To better understand the molecular mechanisms of this process, our colleagues measured fusion rates of a SARS-CoV-2 Omicron containing the H655Y mutation in the spike protein (wt) and a recombinant Y655H revertant^10^. Viral fusion rates were estimated by loading virus-like particles with a luciferase substrate and mixing them with extracellular vesicles containing luciferase. Fusion events are therefore correlated with luciferase activity. Extracellular vesicles and the wt or revertant particles were combined, and luciferase activity was measured every 5 minutes for 3 hours.

The resultant luciferase signal curves are roughly logistic (Fig 4A, Supplemental file 2) but show decay in the “stationary phase” portion of the curve. Fitting a logistic curve to each dataset indicates the revertant has a faster fusion rate (Fig 4B), however goodness-of-fit metrics provided by CurveKinetix indicate this may not be the most appropriate method to derive fusion rates (Fig S2).

**Fig 4.**
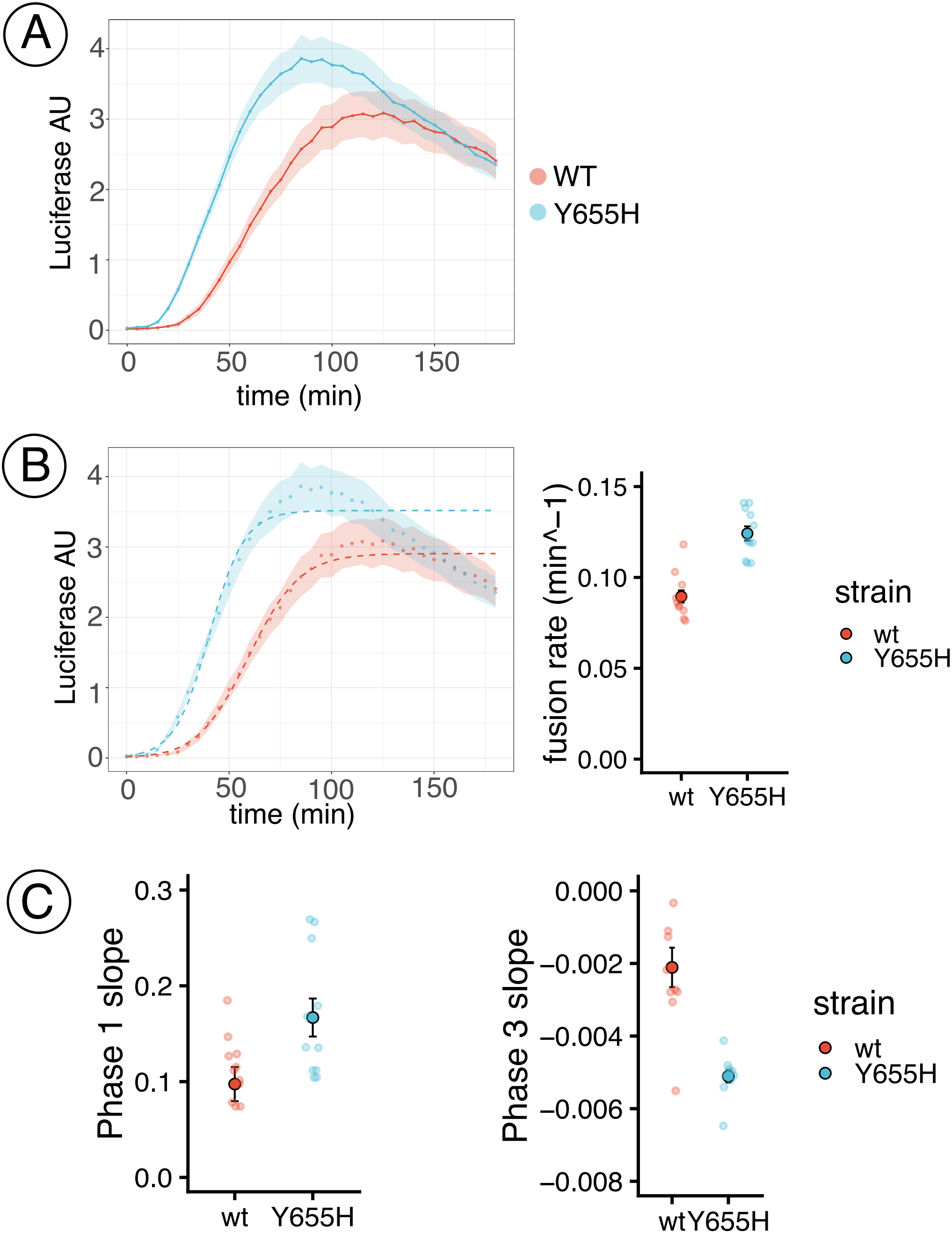
Mutation in the Covid spike protein alters viral fusion dynamics. **A** Luciferase activity overtime during Covid fusion with host cells. A comparison between WT and mutant spike proteins. Line indicates the mean of 3 technical replicates x 3 biological replicates; shaded area is the SEM. **B** A logistic fitting of the luciferase data (left) and the derived fusion rates from this fitting. **C** Luciferase activity during each phase as determined by breakpoint analysis.

We moved to breakpoint analysis to catch the more nuanced shape of the curve to derive more meaningful biological information (Fig 4C,D). Three phases were identified, one with where fusion rates are increasing early in the assay, the second where rates plateau, and the third where rates decrease. From this analysis, the revertant consistently has a higher rate of fusion in the first phase of the assay. However, the revertant also shows a higher rate of luciferase signal decay at the end of the assay, indicating that indeed its fusion is quick and unstable compared to wt, an important observation that was obscured by using a logistic fit to estimate the fusion rate.

The utility of CurveKinetix, therefore, extends outside of the realm of bacterial population dynamics It provides several approaches for summarizing and modeling time-resolved data, making it a general tool for a variety of applications. We can imagine a range of data that CurveKinetix can handle, including fluorescence intensity changes from time-lapse microscopy, fluorescence recovery after photobleaching curves, and enzymatic kinetics.

### Concluding remarks

Biologists routinely measure how variables change over time, but quantitative analysis of these data is not always straightforward. CurveKinetix is a user-friendly application that allows for streamlined time-series data analysis and visualization by people without coding fluency. The hope is that the development and adoption of CurveKinetix may allow for broader use of quantitative analyses, promoting increased tractability, rigor, and stimulation of biological discovery.

## Methods

### Application details and availability

Please see the *Supplemental file* for details on the mathematical transformations used in the data analysis.

CurveKinetix is downloadable as an R package from GitHub ; link to come. Beta version is available at: https://01a165d1-6798-3b46-57e4-5e0b41616ec1.share.connect.posit.cloud/?private_link_token=JcYbUUoqN3Qgn749hAcIbFqlZrLMKOVFBkIsX0uxtDDkHM7tYz8kcqJGwrEqng2T

### Salmonella gene expression case study

Data used for the analysis were generated previously^7^. Briefly, *Salmonella* and *Salmonella* SPI1-*gfp* cells were grown in LB medium in the presence of short-chain fatty acids (acetate, butyrate, and propionate) at different concentrations in a 96-well plate. OD600 and GFP were measured every three minutes. A total of 9 measurements were made; 3 technicals within 3 biological replicates.

The raw plate-reader data were copy-and-pasted into the CurveKinetix manual entry data wizard for each biological replicate, along with the strain and SCFA concentration information for each well. The formatted data from all three biological replicates was merged to create Supplementary File 1.

From there, we moved to background subtraction. The minimum value per curve was subtracted from each OD600 curve; wt GFP fluorescence was subtracted from SPI1-*gfp* GFP fluorescence to account for autofluorescence. From there, we fit logistic curves and performed breakpoint analysis (guided mode, initial guess for breaks at 3 h and 8 h). The resulting data are presented here.

### Viral fusion kinetics

Data used for this analysis were generated previously. The luminescence data were copy-and-pasted into the CurveKinetix manual entry data wizard for each replicate, along with the strain information. The resulting formatted data are in Supplementary File 2.

From there, we fit logistic curves and performed breakpoint analysis (automatic mode, maximum number of breakpoints = 2). The resulting data are presented here.

## Acknowledgements

We are grateful to Crimson Stuckert and Jingwen Zhao for acting as scrupulous beta testers throughout CurveKinetix’s development. and thanks to Enya Qing and Tom Gallagher for sharing the viral fusion dynamics dataset. This work was funded by start-up funds from Loyola University Chicago to Alyson Hockenberry.

## Figure legends

**Fig S1.**
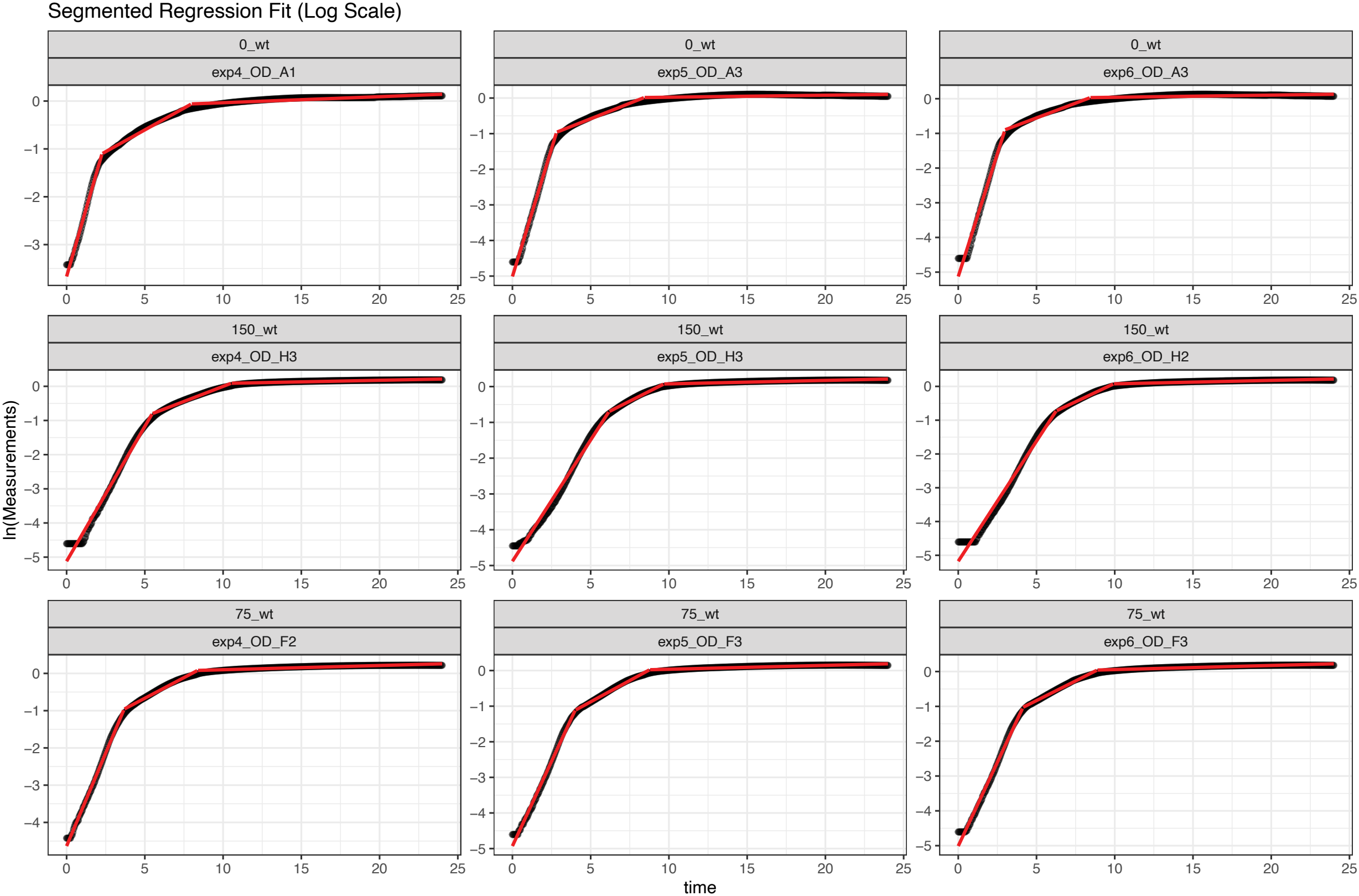
Representative breakpoint analysis outputs for *Salmonella* growth in SCFAs.

**Fig S2.**
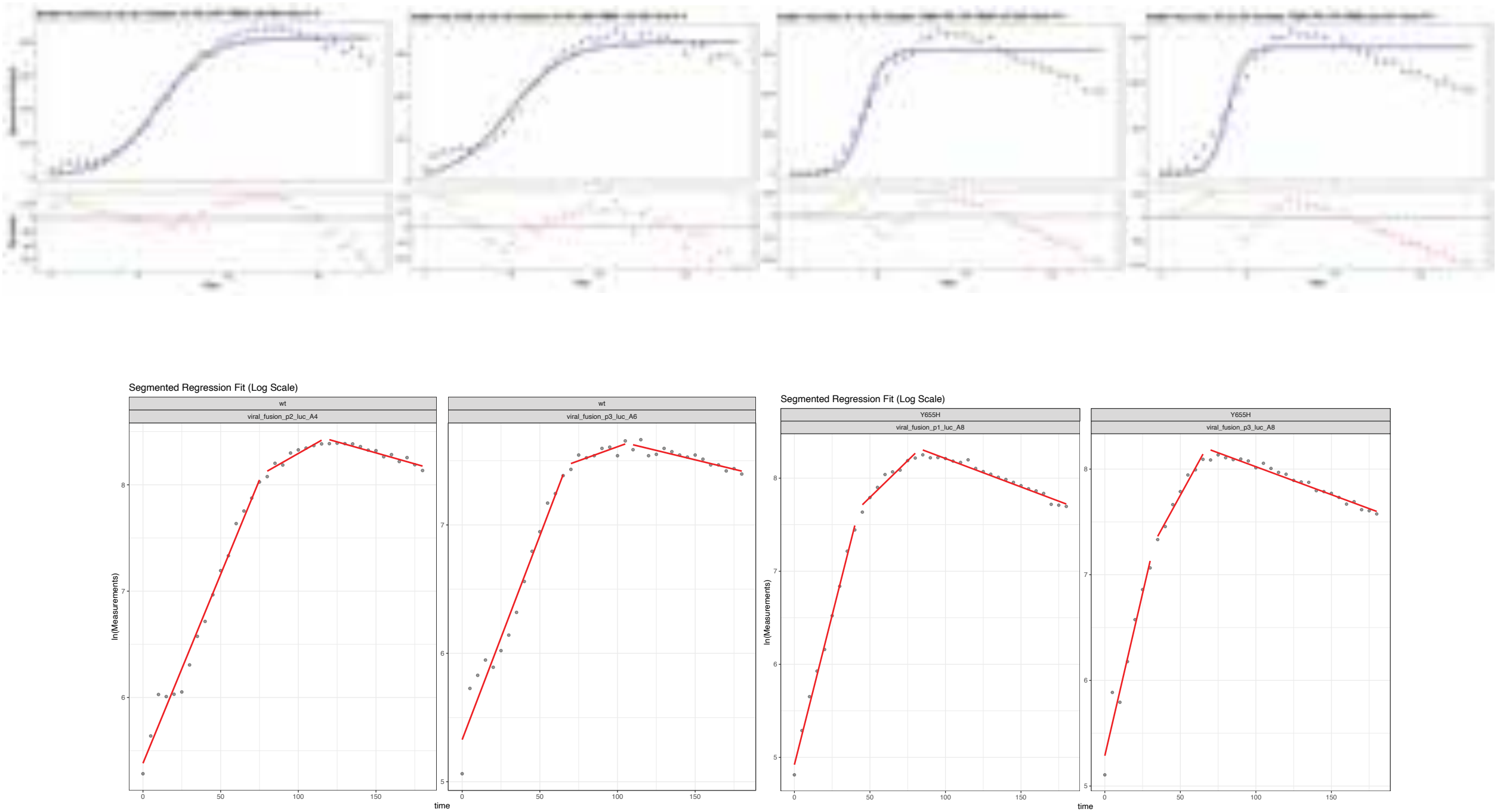
Goodness of logistic vs segmented regression fits in viral fusion analysis.

